# Identification of *Lagopus muta japonica* plant food resources in the Northern Japan Alps using DNA metabarcoding

**DOI:** 10.1101/2021.05.20.444928

**Authors:** Taichi Fujii, Kaoru Ueno, Tomoyasu Shirako, Masatoshi Nakamura, Motoyasu Minami

## Abstract

DNA metabarcoding was employed to identify plant-derived food resources of the Japanese rock ptarmigan (*Lagopus muta japonica*), registered as a natural living monument in Japan, in the Northern Japanese Alps in Toyama Prefecture, Japan, in July to October, 2015-2018. By combined use of *rbcL* and ITS2 local databases of 74 alpine plant species found in the study area, a total of 43 plant taxa were identified and could be assigned to 40 species (93.0%), two genera (4.7%), and one family (2.3%). Rarefaction analysis of each sample collection period showed that this study covered more than 90% of the plant food resources found in the study area. Of the 21 plant families identified using the combined *rbcL* and ITS2 local databases, the most dominant families were Ericaceae (98.1% of 105 fecal samples), followed by Rosaceae (42.9%), Apiaceae (35.2%), and Poaceae (19.0%). In all fecal samples examined, the most frequently encountered plant species were *Vaccinium ovalifolium* var. *ovalifolium* (69.5%), followed by *Empetrum nigrum* var. *japonicum* (68.6%), *Vaccinium* sp. (54.3%), *Kalmia procumbens* (42.9%), and *Tilingia ajanensis* (34.3%). Rarefaction analysis of each collection period in the study revealed that this study covered more than 90% (from 91.0% in July to 97.5% in September) of the plant food resources found in the study area, and 98.1% of the plant food taxa were covered throughout the entire study period. Thus, DNA metabarcoding using the *rbcL* and ITS2 local databases of alpine plants in combination and rarefaction analysis are considered to be well suited for estimating the dominant food plants in the diet of Japanese rock ptarmigans. Further, the local database constructed in this study can be used to survey other areas with similar flora.

## Introduction

The rock ptarmigan (*Lagopus muta*) is a medium-sized grouse that is widely found in subarctic regions of Eurasia and North America [1]. The Japanese rock ptarmigan (*L. m. japonica*), a subspecies of the rock ptarmigan, is endemic to Japan and resides in the southernmost region of the rock ptarmigan’s habitat [1, 2]. The Japanese rock ptarmigan is a relict species that remained on the Japanese archipelago after the last glacial stage. The Japanese rock ptarmigan inhabits the alpine meadow zone, which is beyond the forest limit and scattered with *Pinus pumila*, at altitudes ranging from 2,000 to 3,000 m above sea level (a.s.l.) on the Japanese mainland [1, 3] and feeds mainly on alpine plants found in the region. During the breeding season (April to May), the male occupies its territory of 0.015 to 0.072 km^2^ [3]. The female builds her nest at the base of *P. pumila* from June to July, and lay eggs in July. The female alone incubates the eggs and raises the chicks until October, at which time the chicks become independent [4]. When their habitat (alpine meadow zone) is covered with snow (November to April), the Japanese rock ptarmigan migrates to the forest zone, returning to the alpine meadow zone when the snow begins to melt [5].

Although the Japanese have traditionally protected the Japanese rock ptarmigan as a worship object since ancient times by not allowing their capture or hunting, since the 1930s, populations of this species on several mountains have either declined or become locally extinct [1]. In 1955, the Japanese rock ptarmigan was registered as a natural living monument in Japan because of its worldwide scarcity, and it is the focus of conservation efforts [1]. Haneda et al. [6] estimated the population of Japanese rock ptarmigan to be 3,000 in the 1980s. Despite ongoing conservation efforts, by the early 2000s, the population of Japanese rock ptarmigans was estimated to be 1,700 [1]. The reasons for this decline are poorly understood; however, loss of alpine meadow vegetation and damage by Sika deer (*Cervus nippon*) are thought to be primarily responsible [7]. In addition, the alpine meadow vegetation has been damaged extensively by mountain climbers in the alpine belt [8]. Recently, concerns have been raised about habitat destruction due to climate change. It is predicted that the habitat will be drastically reduced from 2081 to 2100 and that the future risk of extinction is very high [3]. To address these issues, restoration and management of the alpine meadow flora is being carried out in various alpine areas in Japan [9, 10]. However, none of these efforts has taken into consideration the impact of plant foods utilized by the Japanese rock ptarmigan, primarily because few surveys of the plant foods of this species have been conducted to date. In addition, breeding programs have been implemented in an attempt to increase the population; however, chicks raised in captivity are suggested to suffer high mortality as a result of inappropriate dietary composition [11, 12]. For these reasons, detailed understanding of the alpine meadow vegetation as a food source for Japanese rock ptarmigans is important for conservation of this species [13].

Given this background, several methods have been employed to qualitatively or quantitatively elucidate the dietary composition of the Japanese rock ptarmigan. Previously used methods for identifying such plant food resources involved observation of the gastric contents of this species via dissection [14, 15]. However, such destructive methods cannot be employed in studies on the Japanese rock ptarmigan, because of its protected status in Japan. More recently, the most common method for investigating the plant foods of this species is the direct observation of its foraging behavior; however, this method requires specific training in plant species identification and long-term observation by numerous observers [13]. Further, the accuracy of direct observation depends on the observer’s ability to identify plant species, and thus this method is neither objective nor reproducible. In addition, direct observation may cause stress to the animal due to the presence of observers and damage the alpine meadow vegetation by trampling. Another method involves microscopic analysis of plant food residues in feces; however, this method has been shown to be relatively inaccurate. For example, according to a study examining fecal residues in *L. m. pyrenaica* in the French Pyrenees, the plant residues were almost entirely fragmented and could not be identified to the species level [16, 17]. Therefore, it is necessary to establish a stress-free survey method to determine the plant food resources of the Japanese rock ptarmigan at the species level more rapidly than the previously reported survey methods [13–17].

The DNA barcoding method has been successfully applied to food residues in feces and gastric contents to identify dietary composition in wildlife [18–30]. We also reported the first application of DNA barcoding by the cloning method and using the partial sequence of the chloroplast *rbcL* gene (*rbcL*) to plant food residues in the feces of the Japanese rock ptarmigan in Japan’s Northern Alps [31, 32]. Results obtained using a combination of the DNA Data Bank of Japan (DDBJ) and a local database comprised of *rbcL* DNA sequences constructed from 74 alpine plant species found in the study area revealed a total of 26 taxa, 22 species, two genera, and two at the family level [32]. Compared with previous studies of other groups, the DNA barcoding performed using DDBJ and the local database in our previous study [32] was able to identify more plant species than by observation of the gastric contents alone [14, 15], and almost the same number of plant species as direct observation of foraging behavior but in a shorter period [13]. However, the cloning method employed in our previous studies is limited [31, 32].

In the short *rbcL* region used in our previous studies, closely related plant species, e.g., Asteraceae, *Vaccinium* sp., and *Rhododendron* sp., with no polymorphisms in this region have the same homology. Therefore, these taxa could not be identified at the species level. In addition, the rarefaction and extrapolation sampling curves revealed that this survey covered 89% of plant food candidates present in the study area [32]. This means that our previous method may underestimate the number of plant species foraged, as a limited number of DNA sequences were obtained from one fecal sample using the cloning method, and that all plant food candidates in the study area were not detected. Thus, there are major obstacles to identifying plant food residues in feces at the species level and all plant food candidates in the habitat by the cloning method using only the *rbcL* DNA sequences. Recently, DNA metabarcoding using next-generation sequencing (NGS) has been used to identify dietary components in fecal samples of wildlife with higher sensitivity, since this method can determine more DNA sequences than the cloning method [18–30]. Further, plant species identification by DNA barcoding can be improved by combining *rbcL* sequences, which have been deposited for many plants in DNA databanks, and the internal transcribed spacer 2 region between 5.8S ribosomal RNA and 28S ribosomal RNA of nuclear DNA (ITS2) with high interspecific variability [33, 34].

This study aimed to employ a local database of partial sequences of *rbcL* [35] and ITS2 regions [33] for the alpine plants found in the study area (local database), and to identify dietary components by DNA metabarcoding using NGS in reference to the local database. In this study, we used these methods to analyze the major plant food resources of the Japanese rock ptarmigan during the breeding season (July to October) in Japan’s Northern Alps, which is at the center of their distribution area, and the effectiveness of these methods was evaluated. In addition, in order to evaluate the completeness of our sampling efforts, rarefaction analysis was conducted to estimate the percentage coverage of the number of identified plant food taxa relative to the number of analyzed fecal samples [36–39].

## Materials and methods

### Study area

This study was conducted in full compliance with Japanese nature park laws and regulations, including obtaining a license from the Ministry of Environment of Japan to collect Japanese rock ptarmigan feces and plant species. The study was carried out at altitudes ranging from 2,324 to 2,373 m a.s.l. on and around the peak of Mt. Taro (36°26’50.1” N, 137°30’48.9” E, 2373 m a.s.l.), in Chubu-Sangaku National Park at the western end of Japan’s Northern Alps, Toyama Prefecture, Japan (Fig 1). In the study area (ca. 70 ha), the plant community on the northwest-facing slope was dominated by the species *Empetrum nigrum* var. *japonicum*, *Vaccinium uliginosum* var. *japonicum, Sasa kurilensis,* and *P. pumila*; and that on the southeast-facing slope was dominated by the species *Phyllodoce aleutica* and *Kalmia procumbens*. In addition, *Betula ermanii* and family Poaceae grew on the bare field along the wooden footpath, and *Nephrophyllidium crista*-*galli* subsp. *japonicum, Eriophorum vaginatum*, and *Juncus filiformis* grew in and around the small pools.

**Fig 1.**
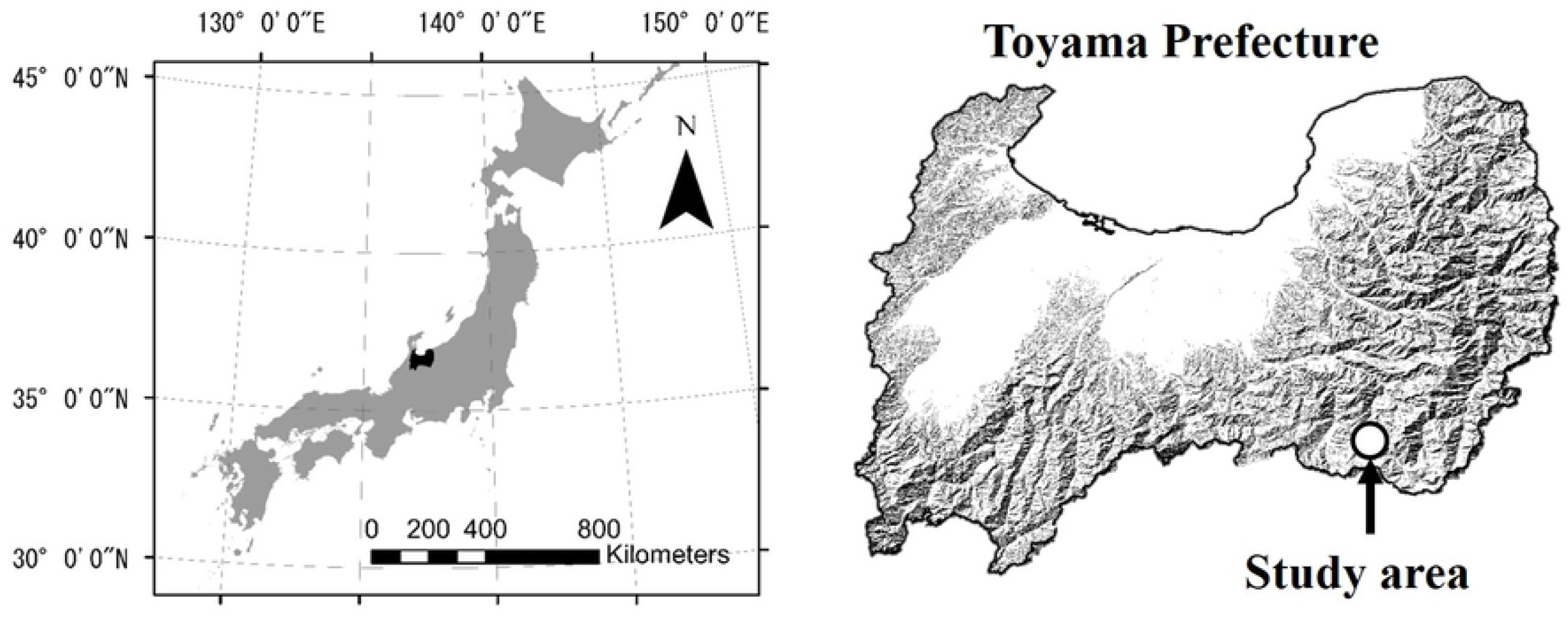
Location of the study area on Mt. Taro in Chubu-Sangaku National Park, at the western edge of the Northern Alps, Toyama Prefecture, Japan.

### Collection of fecal samples and plants

In the study area, typically three pairs of Japanese rock ptarmigans are observed every year. Since Japanese rock ptarmigans show no fear of humans, continuous monitoring is possible for collection of fecal samples. All fecal samples were collected while mainly monitoring the covey, consisting of the adult female and her chicks, from July to October, 2015-2018 (Table 1). The female is always in the presence of her chicks, and, according to our observations, the plant species foraged by the female and her chicks were almost the same. Therefore, the feces of the female and her chicks were not considered separately, and a total of 117 fecal samples were collected. Samples of 74 alpine plant species were collected from the study area on July 27, 2016 (Table 2) and used to construct the local database of *rbcL* and ITS2 sequences.

**Table 1.** Fecal sampling collection date, number of fecal samples used in the experiment, and number of fecal samples successfully DNA metabarcoding.

| Feces sampling | Number of collected fecal samples | Number of fecal samples successfully amplified both of *rbcL* and ITS2 |
| --- | --- | --- |
| July 8-10, 2016 | 33 | 31 |
| July 26-28, 2016 | 18 | 18 |
| August 18-20, 2015 | 33 | 31 |
| September 13-14, 2018 | 20 | 17 |
| October 11-12, 2016 | 13 | 8 |
| Total | 117 | 105 |

**Table 2.**
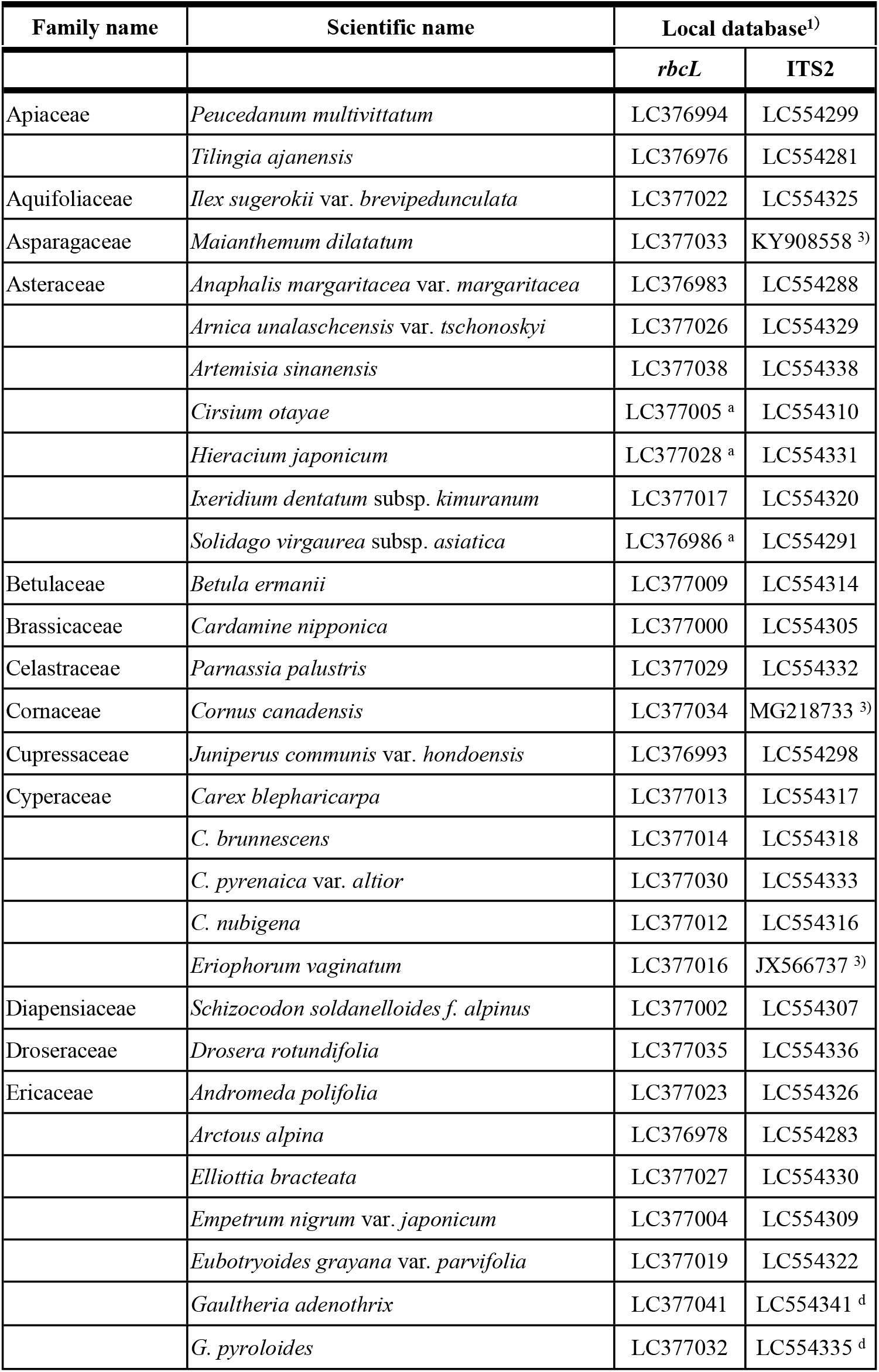
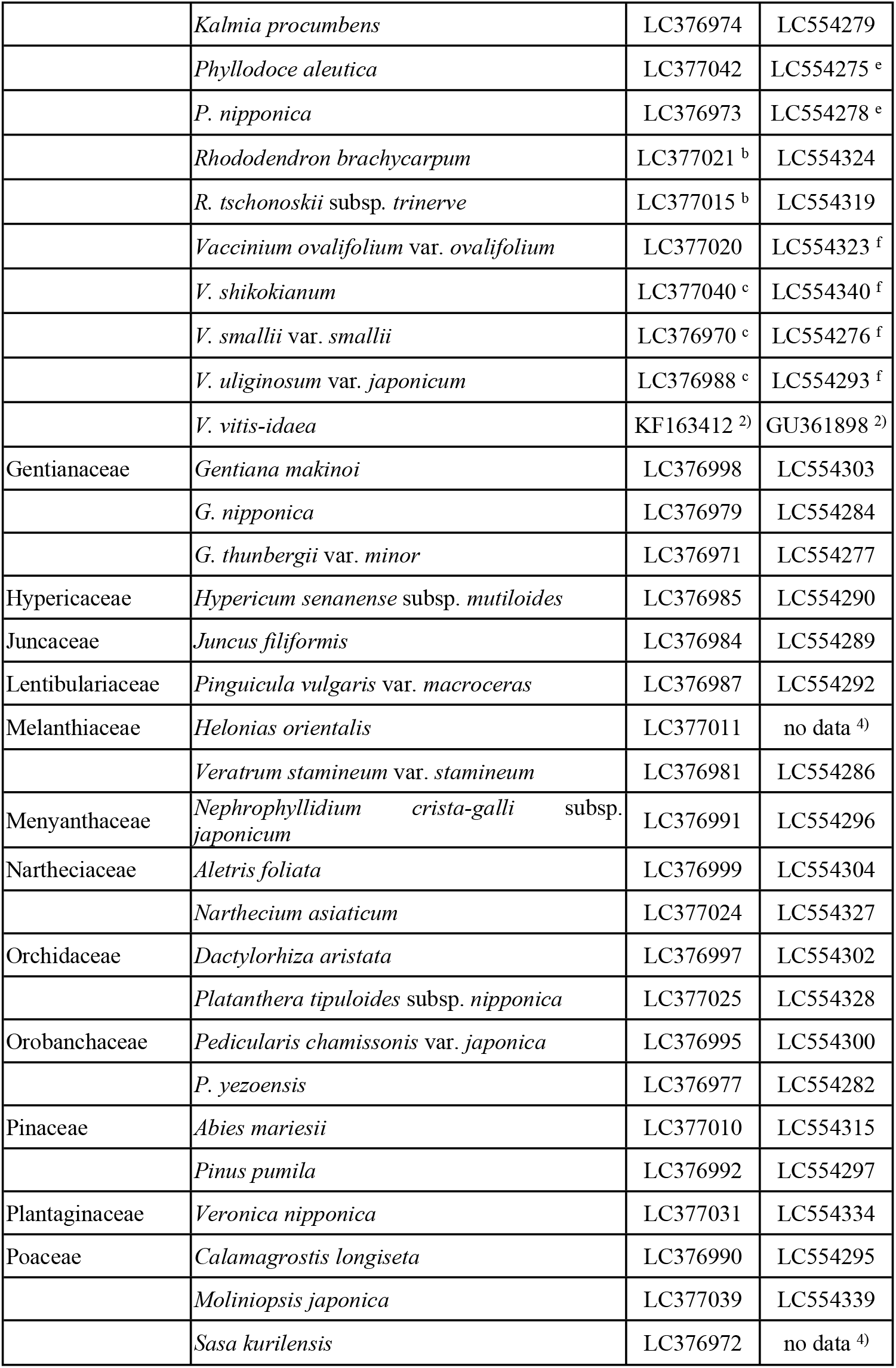
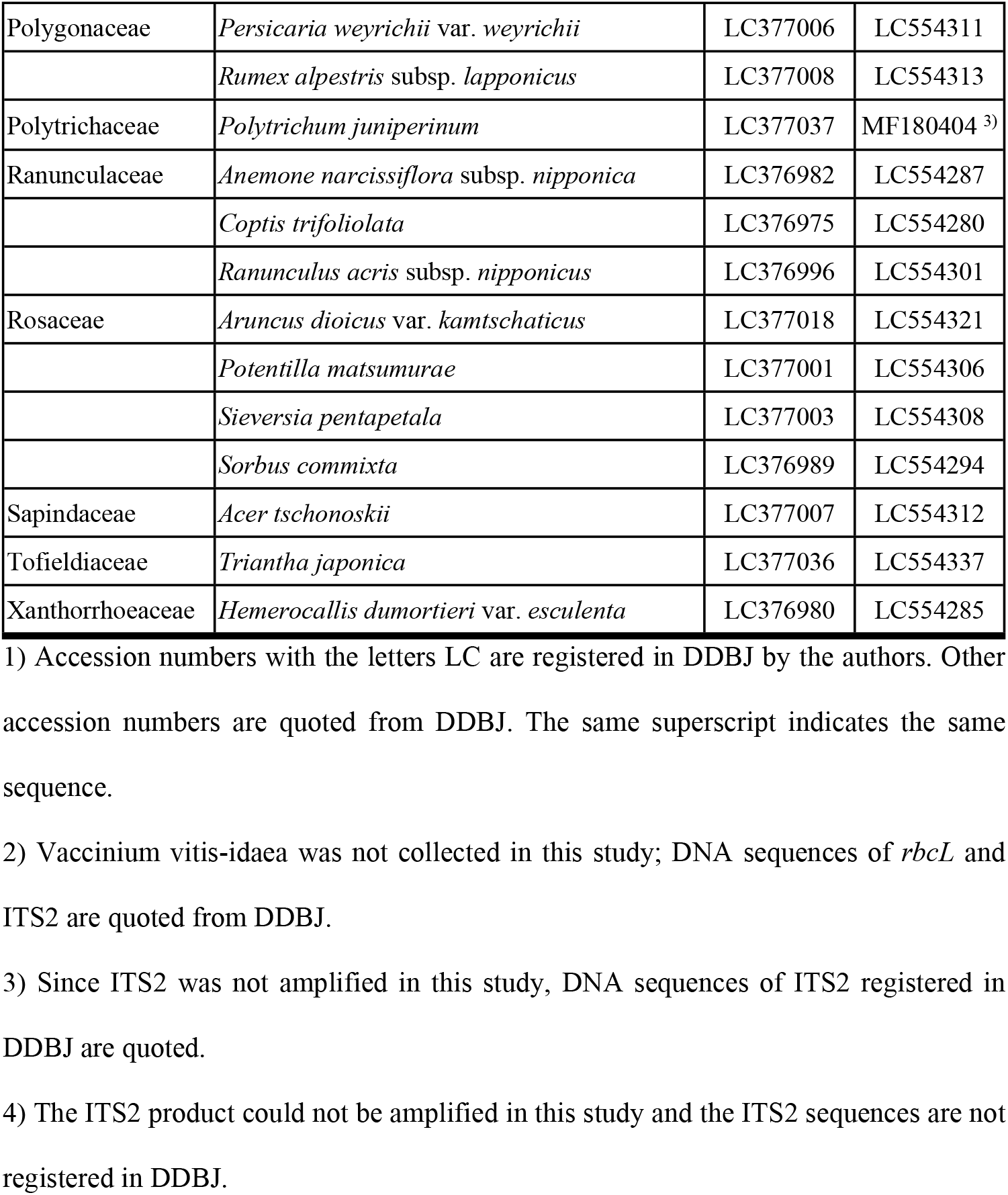
Local databases constructed from alpine plants in the study area.

### DNA metabarcoding of chloroplast *rbcL* and nuclear ITS2

Total DNA was isolated from the dried feces (ca. 2.9-59.2 mg) using a DNeasy Plant Mini Kit (Qiagen, Hilden, Germany) and purified using a Geneclean Spin Kit (MP-Biomedicals, Santa Ana, CA, USA). In this study, DNA metabarcoding using NGS to identify plant species in feces was conducted using the partial sequences of *rbcL* [35] and ITS2 [33]. Since amplicon sequencing by NGS generally decreases nucleotide diversity, adjacent DNA sequences are misrecognized on the flow cell and sequencing quality scores are reduced [40]. Therefore, in order to increase the nucleotide diversity, we used frame-shifting primers [40] for the initial PCR (Table 3). The initial PCRs of *rbcL* and ITS2 were performed in reaction mixtures of 12.0 μl with 6.0 μl of KAPA HiFi (Kapa Biosystems, Wilmington, MA, USA), 0.7 μl of 10 μM primer mix for *rbcL* and ITS2 each (Table 3), 2.0 μl of template DNA, and 3.3 μl of dH_2_O. The initial PCR of *rbcL* was under the following conditions: initial denaturation at 95°C for 3 min, followed by 35 cycles of denaturation at 98°C for 20 s, annealing at 56°C for 15 s, and extension at 72°C for 30 s, with final extension at 72°C for 5 min. The initial PCR of ITS2 was performed under the same conditions as that of *rbcL,* except that the annealing temperature was 55°C instead of 56°C. The initial PCR products of *rbcL* and ITS2 were purified with an Agencourt AMPure XP kit (Beckman Coulter, Fullerton, CA, USA). To construct the DNA libraries using the second PCR, the i7 and i5 indexes (Illumina, San Diego, CA, USA) for identifying each fecal sample and P5 and P7 adapters (Illumina) for Miseq sequencing were ligated to the purified initial PCR products. The second PCRs of *rbcL* and ITS2 were performed in reaction mixtures of 24.0 μl with 12.0 μl of KAPA HiFi (Kapa Biosystems), 2.8 μl of 10 μM forward and reverse index primers (Table 4), 2.0 μl of 1/10 1st PCR products, and 4.4 μl of dH_2_O. The second PCRs of *rbcL* and ITS2 were under the following conditions: initial denaturation at 95°C for 3 min, followed by 12 cycles of denaturation at 98°C for 20 s and extension at 72°C for 15 s, with final extension at 72°C for 5 min. The DNA libraries of *rbcL* and ITS2 purified with an Agencourt AMPure XP kit (Beckman Coulter) were sequenced using a MiSeq Reagent Kit v3 (600-cycle format; Illumina) with the Illumina MiSeq sequencer following the manufacturer’s protocol.

**Table 3.**
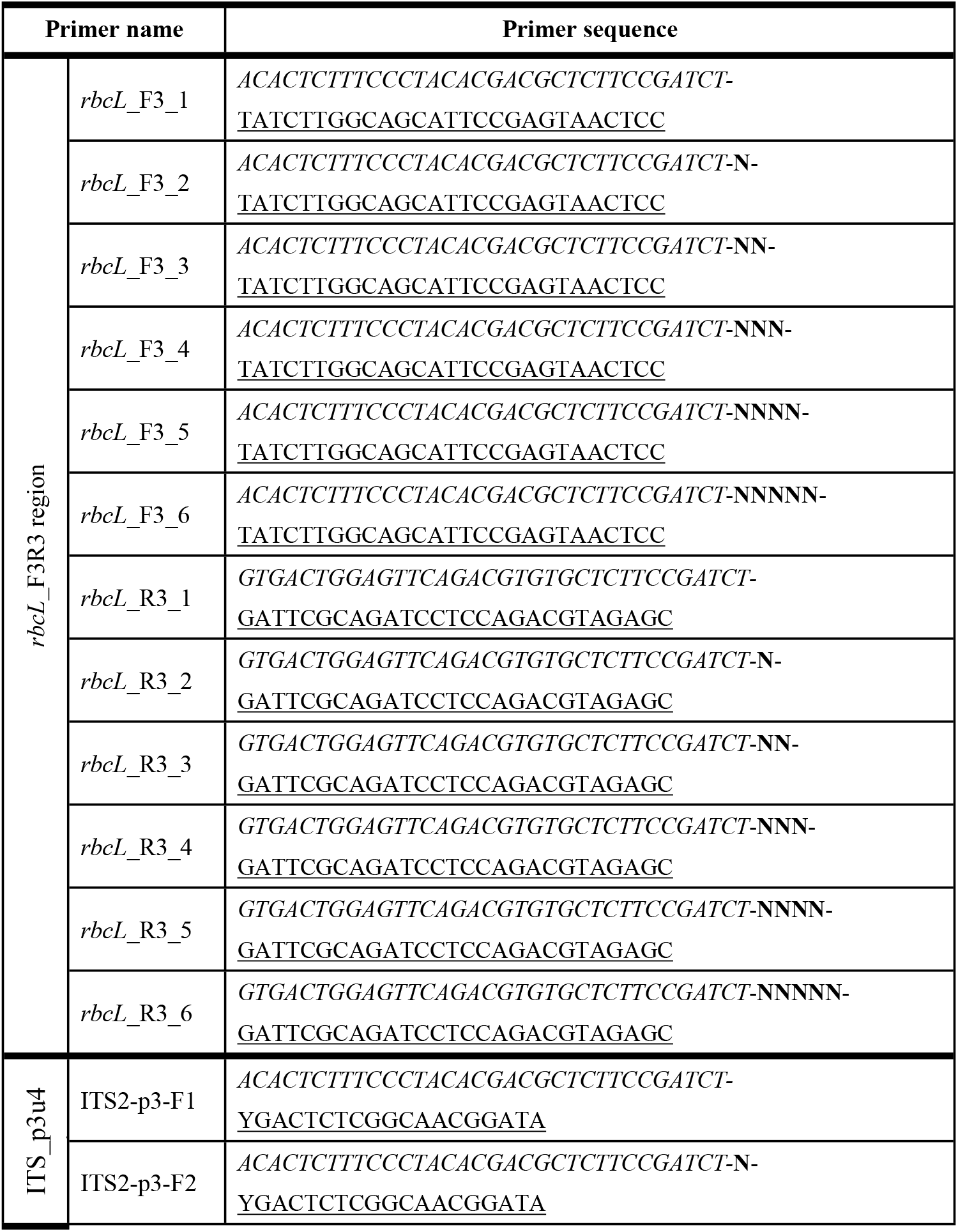
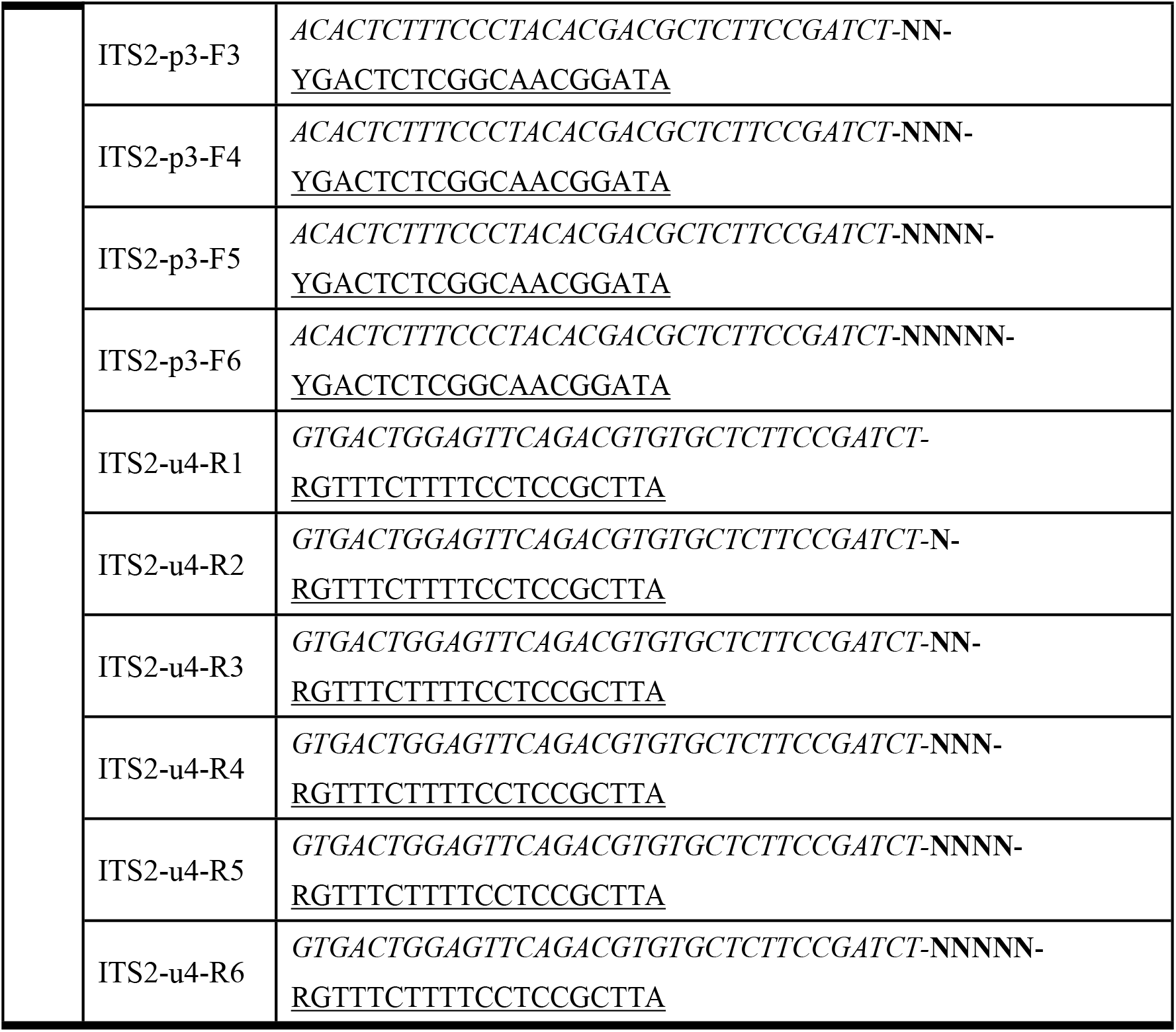
List of initial PCR primers.

**Table 4.**
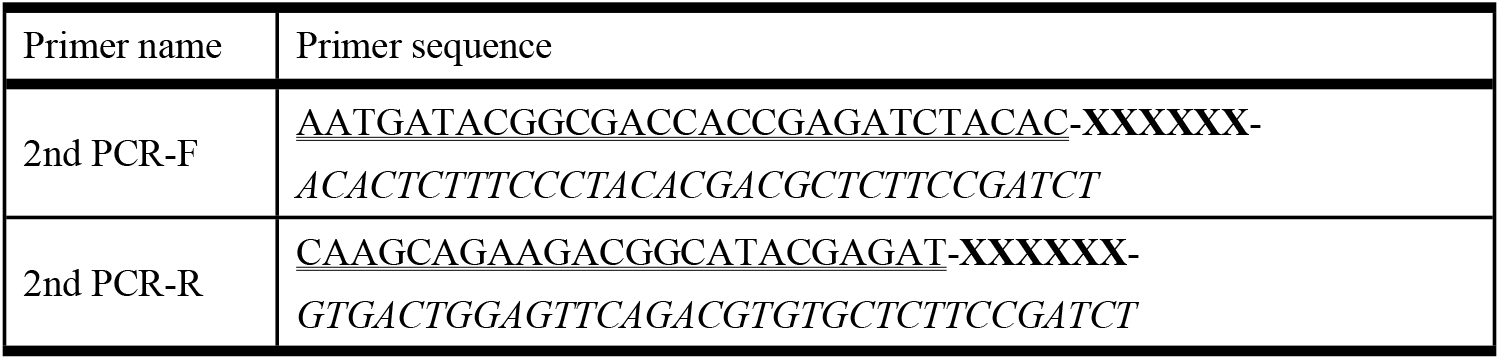
List of second PCR primers.

Italic characters indicate MiSeq sequencing primers. Bold Ns indicate random bases to improve the quality of MiSeq sequencing [40]. Single underlined characters indicate DNA barcoding primer sequence.

Italic characters indicate MiSeq sequencing primers. Bold Xs indicate index sequences to identify each sample. Double underlined characters indicate P5/P7 adapter sequences for capture to flow cell.

### Sequence data analysis

Since the demultiplexing function in the Illumina MiSeq sequencer (Illumina) does not evaluate the sequencing quality scores of the i7 and i5 index sequences, the default FASTQ files provided by the Illumina MiSeq sequencer (Illumina) often contain miss-tagged sequences [41]. To solve this problem, the raw MiSeq data were converted into FASTQ files using the bcl2fastq v2.18 program (Illumina), and the FASTQ files were then demultiplexed using the clsplitseq function implemented in Claident [42] instead of the default demultiplexing function in the Illumina MiSeq sequencer (Illumina). In general, demultiplexed FASTQ files generated by the MiSeq sequencer are analyzed using either the ASV (amplicon sequence variant) or OTU (operational taxonomic unit) approach [43]. In this study, the demultiplexed FASTQ files were analyzed using the ASV method implemented in the DADA2 v1.10.1 package [43], with the statistical software R [44]. ASV can discriminate between DNA sequences that differ by a single nucleotide, since the original DNA sequence can be estimated even if it contains PCR amplification or sequencing errors [43]. At the quality filtering process, forward and reverse sequences of both *rbcL* and ITS2 sequences were trimmed at 200 and 250 bases, respectively, based on visual inspection of the quality score distribution, using the filterAndTrim function of DADA2 [43]. After trimming, forward and reverse sequences were combined using the mergePairs function of DADA2 [43], and chimeric DNA sequences were removed using the removeBimeraDenovo function of DADA2 [43]. Low-frequency ASVs, i.e., less than 1.0% of the total number of sequences, in each fecal sample were excluded from the DNA metabarcoding analysis. To normalize the number of DNA sequences in each *rbcL* and ITS2, rarefaction curves were calculated with the rarecurve function of vegan package ver.2.5-5 [45] in the statistical software R [44]. Using the calculated rarefaction curves, 1,000 DNA sequences in each *rbcL* and ITS2 were normalized for each fecal sample using the rrarefy function of the vegan R package [45] to determine the ASVs for plant species identification.

### Construction of *rbcL* and ITS2 local databases for the fecal samples collection area

In this study, we used the local database of *rbcL* from our previous study [32] (Table 2). Since the *rbcL* sequence is haploid, its DNA sequence can be determined by the direct sequencing method. On the other hand, since ITS2 is nuclear DNA, it may contain heterogeneous sequences within the same plant species. Therefore, the DNA sequencing for ITS2 was performed by the amplicon sequence method using NGS. Using the same DNA extracts for the construction of the local *rbcL* database [32], PCR amplification of ITS2 was performed using the same reaction mixture and conditions as the initial and second PCRs of ITS2. The PCR products of ITS2 purified with an Agencourt AMPure XP kit (Beckman Coulter) were subjected to MiSeq sequencing. When the same plant species had multiple DNA sequences of ITS2, the DNA sequence with the highest number of reads was adopted as the DNA sequence for the ITS2 local database and was registered in DDBJ (for detailed DDBJ accession nos. of each *rbcL* and ITS2, refer to Table 2).

### Homology search and identification of plant food resources

In this study, only fecal samples in which both *rbcL* and ITS2 could be PCR amplified were used as food plant identification samples. To identify food plant species, homology searches were performed by comparing the ASVs obtained for each fecal sample against the DNA sequences of *rbcL* and ITS2 in our local database using OmicsBox ver. 1.4 [46]. To ensure that the identification accuracy was sufficiently robust, plant species with a homology of less than 98% were excluded from further analysis [31, 32]. In the event that a given sequence was identified as belonging to two or more taxa with the same score, that sequence was assigned to the highest taxonomic level that included both of those taxa [31, 32]. As a result, some of the obtained DNA sequences were assigned to the rank of genus and others to family. Eventually, we compared the taxon level identified by *rbcL* or ITS2, and selected the lower taxonomic level. The species estimated accuracy was calculated by dividing the number of plant taxa of estimated to the species level by the number of total plant taxa.

### Statistical analysis

The numbers of plant taxa per fecal sample identified by the *rbcL* local database, the ITS2 local database, and a combination of the *rbcL* and ITS2 local databases were compared by the Steel-Dwass test (P<0.05) using the pSDCFlig function of the NSM3 package of R [47]. In addition, we calculated the percentage coverage as the number of identified food plant taxa relative to the number of analyzed fecal samples and estimated asymptotic Shannon diversity index by rarefaction analysis using the iNEXT package of R [38, 39, 48].

## Results

### Local database construction

Seventy-three alpine plant specimens, consisting of 32 families and 61 genera, were collected from the study area (Table 2) and used to construct the local *rbcL* and ITS2 databases. Although *rbcL* had already been successfully amplified from all 74 plant specimens in our previous study (Table 2), some taxa could only be identified at the genus or family level due to same *rbcL* sequences; three species in family Asteraceae (*Cirsium <u>otayae</u>, Hieracium japonicum,* and *Solidago virgaurea* subsp. *asiatica*), two species in genus *Rhododendron* (*R. brachycarpum* and *R. tschonoskii* subsp. *trinerve*), and three species in genus *Vaccinium (V. shikokianum, V. smallii* var. *smallii,* and *V. uliginosum* var. *japonicum*) [32]. On the other hand, the ITS2 sequences could be amplified from 67/74 plant specimens and the ITS2 sequences were deposited in DDBJ under accession nos. LC554275-LC554341 (Table 2). For the four out of six ITS2 sequences that could not be amplified in this study, sequences used in our local database of ITS2 sequences were obtained from DDBJ (*Cornus canadensis,* MG218733, DDBJ accession no.; *E. vaginatum,* JX566737; *Maianthemum dilatatum,* KY908558; and *Polytrichum juniperinum*, MF180404). When referencing ITS2 sequences of these four species from DDBJ, ITS2 pseudogene sequences and sequences containing Ns (unknown nucleotides) were excluded from our analysis to ensure identification accuracy. The ITS2 of the remaining two plant species (*Helonias orientalis* and *S. kurilensis*) were not registered in DDBJ. In addition, two species in genus *Gaultheria (G. adenothrix* and *G. pyroloides*), two species in genus *Pyllodoce* (*P. aleutica* and *P. nipponica*), and four species in genus *Vaccinium (V. ovalifolium* var. *ovalifolium, V. shikokianum, V. smallii* var. *smallii,* and *V*. *uliginosum* var. *japonicum*) had the same ITS2 sequences. Our previous study showed that *Vaccinium vitis-idaea* was identified from the fecal samples of Japanese rock ptarmigans in our study area [31, 32]; however, we were unable to collect this plant species in isolation. Therefore, we adopted the *rbcL* (KF163412) and ITS2 (GU361898) sequences of this plant species registered in DDBJ (Table 2). Finally, by combining the *rbcL* and ITS2 local databases, we constructed a local database of 71 species and one genus (genus *Vaccinium*: including *V. shikokianum*, *V. smallii* var. *smallii,* and *V. uliginosum* var*. japonicum*) (Table 2).

### Sequence data processing

Of the 117 fecal samples, both *rbcL* and ITS2 were successfully amplified in 105 fecal samples (Table 1). Ninety-two and 261 ASVs were distinguished from each 105,000 DNA sequence of *rbcL* and ITS, respectively (S1, S2 Tables). Based on the results of homology search using our *rbcL* and ITS2 local databases, the number of ASVs with more than 98% homology were 66 (71.7%) out of 92 ASVs for *rbcL* and 153 (58.6%) out of 261 ASVs for ITS2, respectively. To verify the accuracy of our *rbcL* and ITS2 local databases, the 26 ASVs (28.3%) in *rbcL* and 108 ASVs (41.4%) in ITS2, which had less than 98% homology using our local database, were subjected to homology search using DDBJ. Sixteen out of 26 ASVs for *rbcL* were identified as plant species with more than 98% homology, but not all identified plant species grew in the alpine meadow zone of Japan; the remaining 10 ASVs had less than 98% homology (S3 Table). On the other hand, 44 out of 110 ASVs for ITS2 were identified as plant species with more than 98% homology. Of 44 ASVs, 29 ASVs were identified as *V. uliginosum,* which grows in the study area, while 15 ASVs with more than 98% homology did not grow in the alpine meadow zone and the remaining 66 ASVs had less than 98% homology (S4 Table). Our local database was shown to cover the plant food candidates utilized by the Japanese rock ptarmigan in the study area. By using our local database, the accuracy of plant species identification was higher than that which could be achieved by DNA barcoding using DDBJ alone.

### Identification of plant food resources

The *rbcL* local database results showed 66 ASVs with more than 98% homology, which could be assigned to 36 plant taxa; 34 (94.4%) to species, one (2.8%) to genus, and one (2.8%) to family (S5 Table). On the other hand, the results obtained using the ITS2 local database revealed 153 ASVs with more than 98% homology, which could be assigned to 29 plant taxa; 26 (89.7%) to species, and three (10.3%) to genus (S6 Table). By using the *rbcL* and ITS2 local databases in combination, a total of 43 plant taxa were identified and could be assigned to 40 species (93.0%), two genera (4.7%) and one family (2.3%) (Table 5). The significantly highest number of identified plant taxa per fecal sample was found in the results of *rbcL* and ITS2 local databases in combination (median 5, interquartile ranges 3–6), followed by the *rbcL* local database (4, 3–5), and the ITS2 local database (3, 2–4), (Steel-Dwass test, P<0.05) (Fig 2).

**Fig 2.**
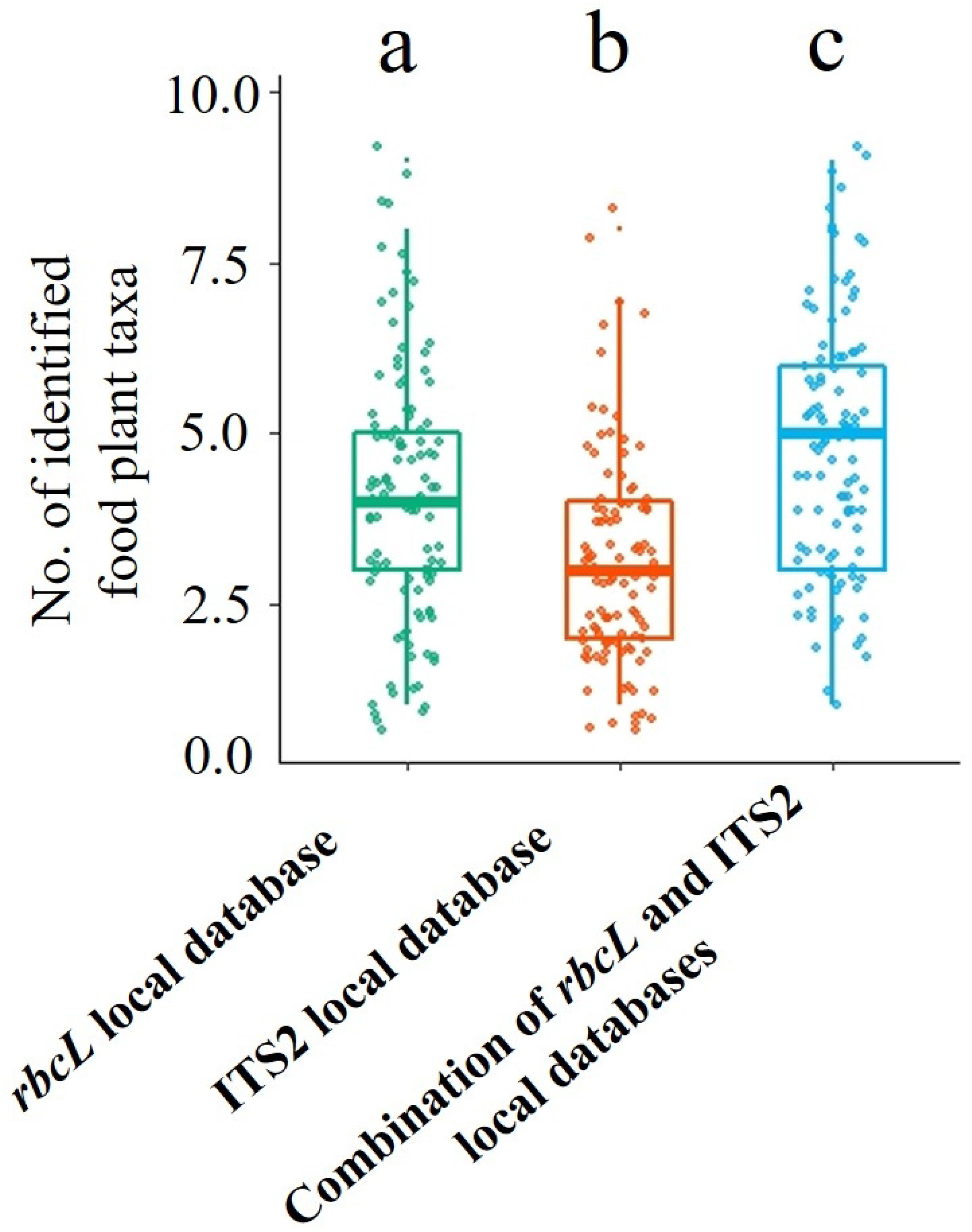
Comparison of the number of plant taxa per one fecal sample identified by the *rbcL* local database, ITS2 local database, and a combination of the *rbcL* and ITS2 local databases. The number of plant taxa per one fecal sample identified by the *rbcL* local database, ITS2 local database, and a combination of the *rbcL* and ITS2 local databases are presented as boxplots. Each box delimits values between 25% and 75% of the group. The bold horizontal line represents the median of the group. Whiskers are drawn for obtained values that differ least from the median ± 1.5 interquartile ranges. Different letters show statistically significant differences (Steel-Dwass test, P<0.05) between groups.

**Table 5.**
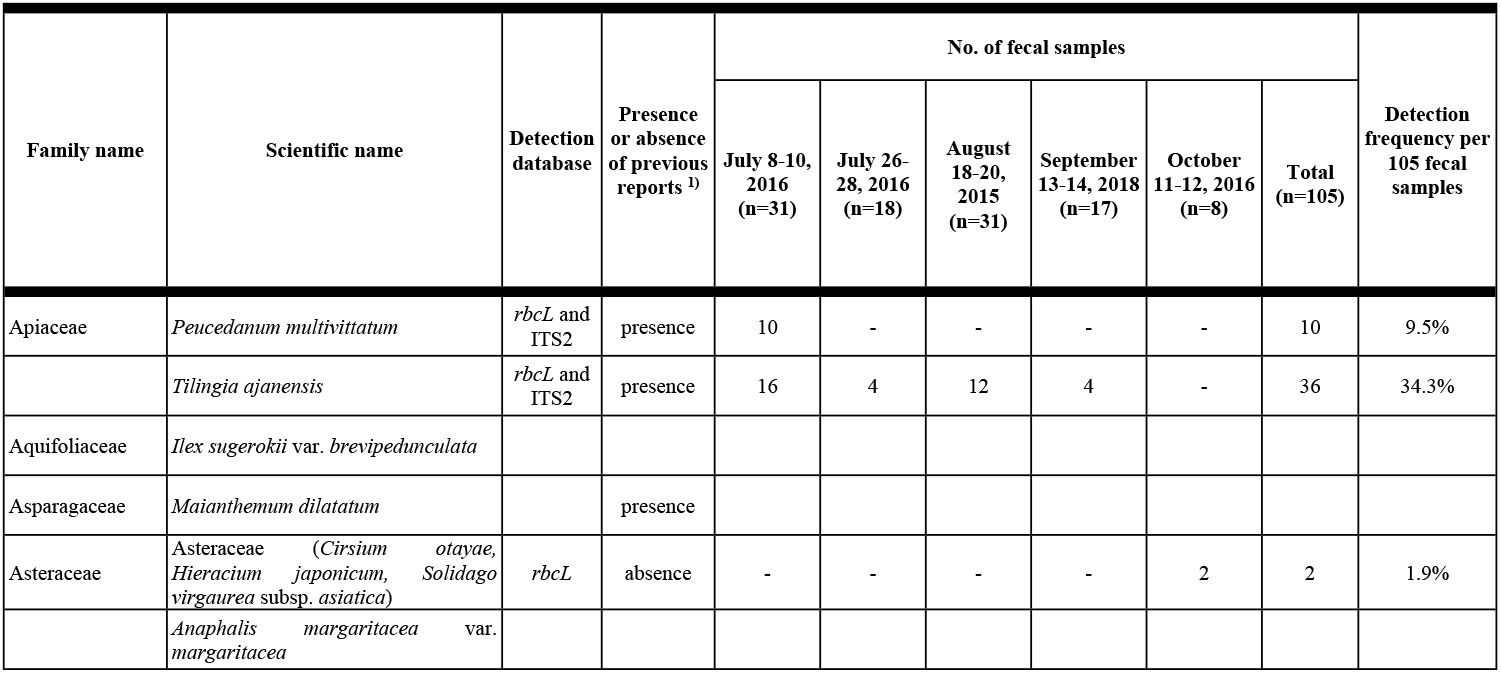
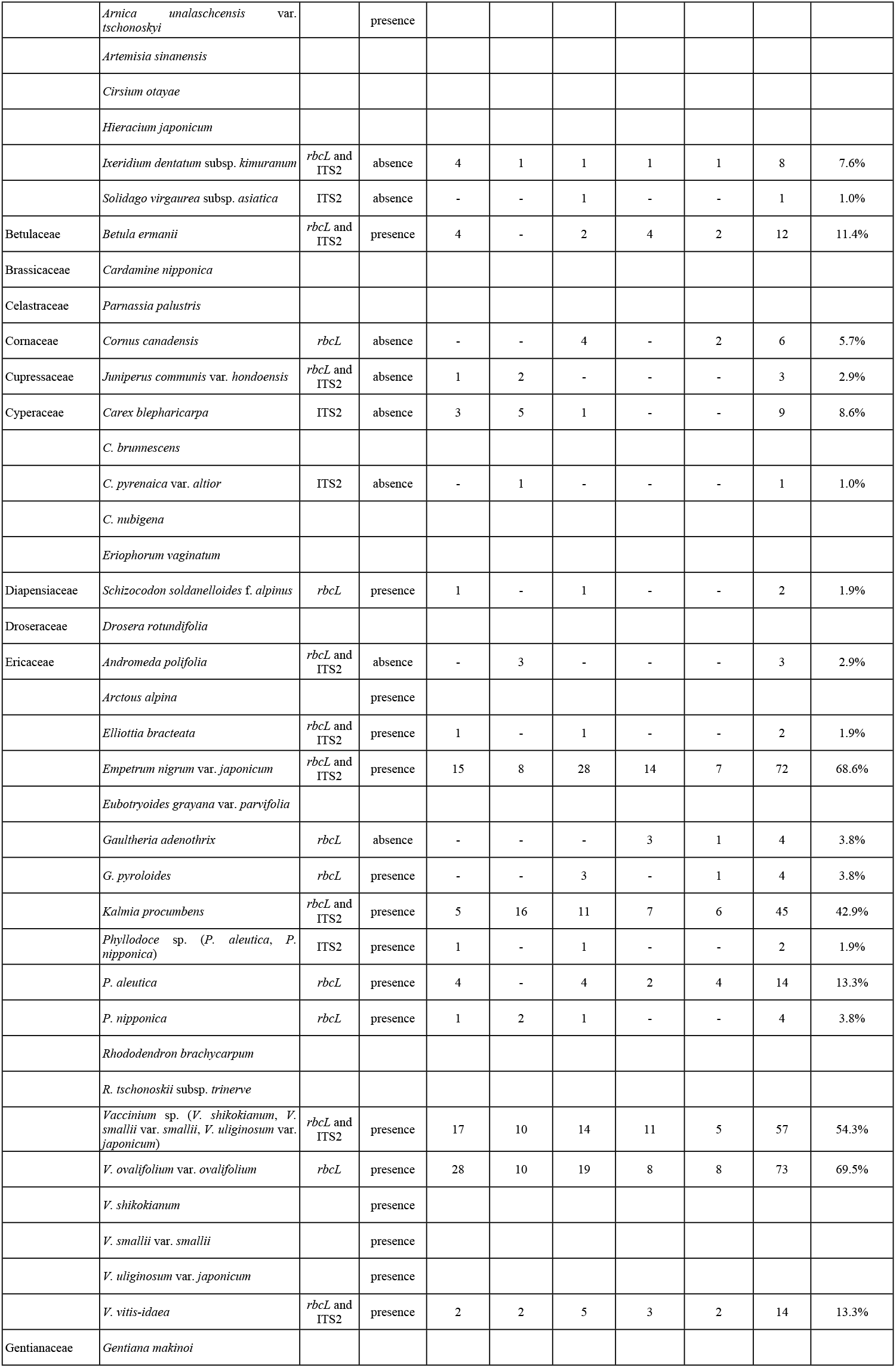
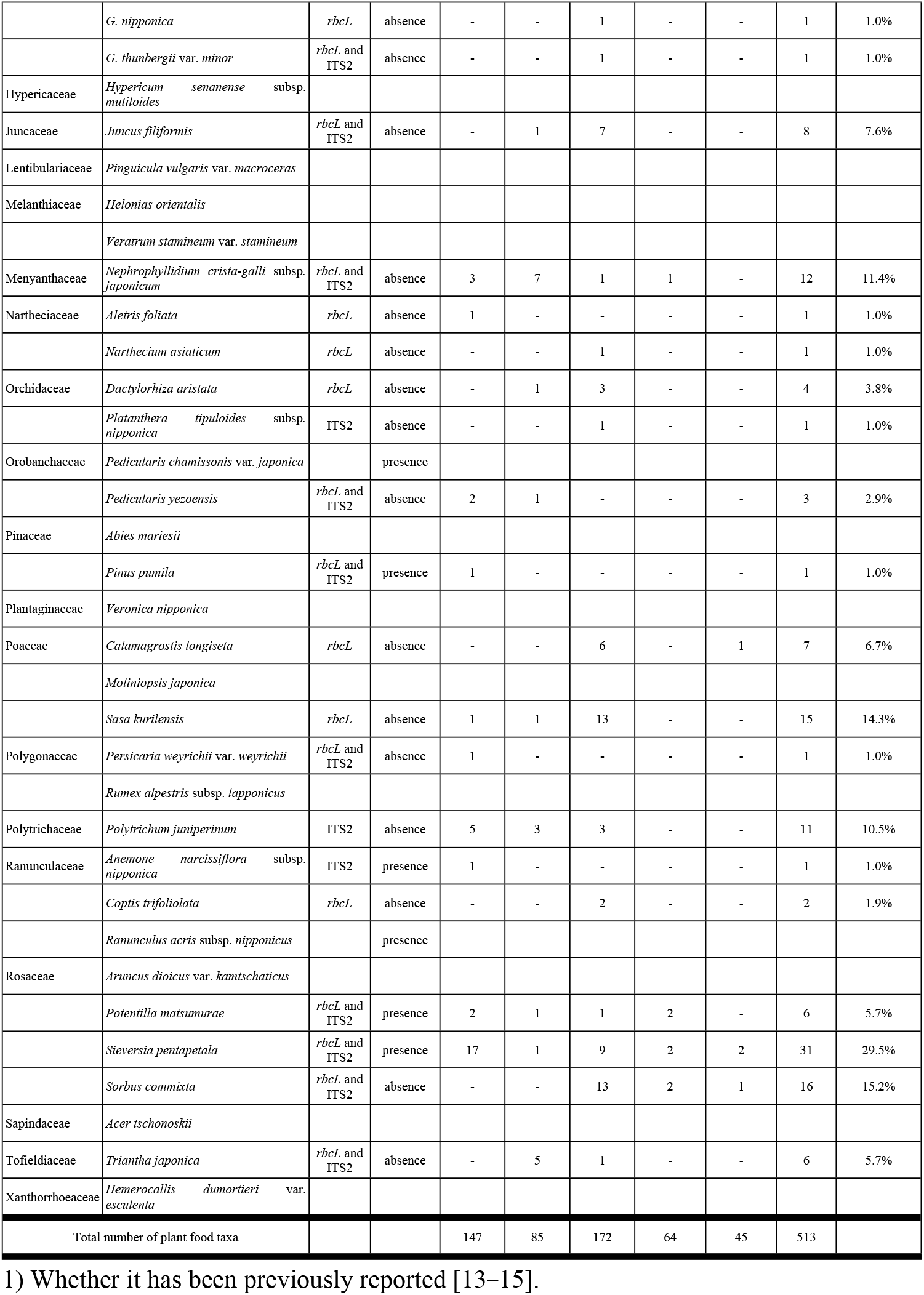
Identified food plant taxa using DNA metabarcoding with *rbcL* and ITS2.

Of the 21 plant families identified using the *rbcL* and ITS2 local databases together, the dominant families throughout all collection periods were Ericaceae (98.1% of 105 fecal samples), followed by Rosaceae (42.9%), Apiaceae (35.2%), and Poaceae (19.0%). In all of the fecal samples examined, the dominant plant foods (more than 30% of 105 fecal samples) were *V. ovalifolium* var. *ovalifolium* (69.5%), followed by *E. nigrum* var. *japonicum* (68.6%), *Vaccinium* sp. (54.3%), *K. procumbens* (42.9%), and *Tilingia ajanensis* (34.3%) (Table 5). Remarkably, *V. ovalifolium* var. *ovalifolium, E. nigrum* var. *japonicum,* and *Vaccinium* sp. were foraged throughout the study period, indicating that these three plant taxa were the most important food resources in the study area (Fig 3). The subdominant plants (more than 10% to less than 30% of 105 fecal samples) were *Sieversia pentapetala* (29.5%), *Sorbus commixta* (15.2%), *S. kurilensis* (14.3%), *P. aleutica* (13.3%), *V. vitis-idaea* (13.3%), *B. ermanii* (11.4%), *N. crista-galli* subsp. *japonicum* (11.4%) and *P. juniperinum* (10.5%) (Table 5). Remarkably, of the subdominant plants, there was abundant foraging of *S. commixta* and *S. kurilensis* in August (Fig 3 and Table 5). The estimated asymptotic Shannon index diversity varied over the study period; the diversity of foraged plant taxa tended to be higher from July to August (16.6–22.1) than September to October (11.3–13.8) (Table 6). Rarefaction analysis of each collection period in the study revealed that this study covered more than 90% (from 91.0% in July to 97.5% in September) of the plant food resources found in the study area, and 98.1% of the plant food taxa were covered throughout the entire study period (Table 6).

**Fig 3.**
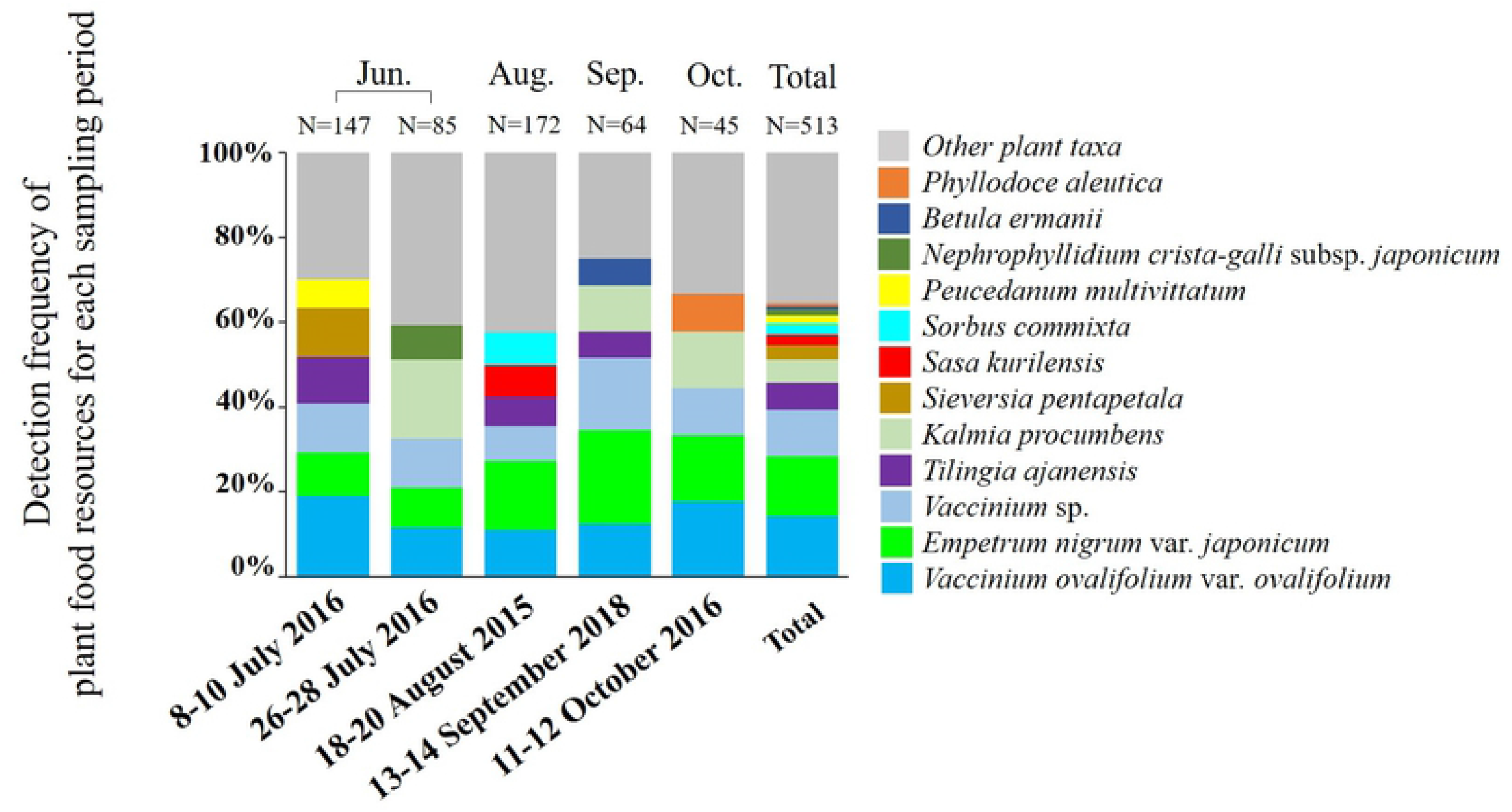
Intra-annual changes of major plant food taxa. The predominant plant species for each of the five fecal sampling periods are shown. N shows the total number of plant food taxa for each fecal sampling period.

**Table 6.** Estimated asymptotic Shannon diversity and coverage of food plant candidates for each fecal sampling period.

| Sample collection period | No. of fecal samples | No. of identified plant taxa | S<sub>obs</sub> | S<sub>est</sub> | Coverage of food plant candidates (%) |
| --- | --- | --- | --- | --- | --- |
| July 8-10, 2016 | 31 | 26 | 14.4 | 16.6 | 93.3 |
| July 26-28, 2016 | 18 | 21 | 14.0 | 16.7 | 91.0 |
| August 18-20, 2015 | 31 | 32 | 18.3 | 22.1 | 91.9 |
| September 13-14, 2018 | 17 | 14 | 10.3 | 11.3 | 97.5 |
| October 11-12, 2016 | 8 | 15 | 11.5 | 13.8 | 91.4 |
| Total | 105 | 43 | 20.0 | 21.1 | 98.1 |

## Discussion

In this study, a total of 43 plant taxa were identified, which could be assigned to 40 species, two genera, and one family. Further, rarefaction analysis of each fecal sample collection period showed that 98.1% of plant food taxa in the study area were covered throughout the entire study period. In a previous study of the gastric contents of the Japanese rock ptarmigan, 17 plant species were identified from 39 individuals in July to October 1926-1928 [14], and only five plant species were observed in the gastric contents of an adult male in October 1967 [15]. Japanese rock ptarmigans inhabiting Mt. Norikura of Japan’s Northern Alps were observed to forage 34 plant taxa, by direct observation of foraging behavior over 43 days from July to October 2009 [13]. Compared with these previous studies, DNA metabarcoding using the local database in this study was able to identify more plant species than was possible using observations of gastric contents [14, 15] and direct observations of foraging behavior [13]. Of the 40 species identified in this study, 24 species and one taxon (identified as family Asteraceae, and either *C. <u>otayae</u>, H. japonicum*, or *S. virgaurea* subsp. *asiatica*) not previously reported as plant food from July to October were identified for the first time in our study [13, 14, 15] (Table 5). In addition, compared with previous studies on the identification of plant foods of other organisms by DNA metabarcoding, the accuracy of this study was high (Table 7). ITS2 is generally considered to be more accurate in diet identification by DNA barcoding than chloroplast DNA [49], because ITS2 has higher interspecific variation than *rbcL* [50, 51]. However, in the 74 alpine plant species used to construct the local database in this study, the percentage of interspecific DNA sequence variation did not differ between *rbc*L (89.2%) and ITS2 (88.9%). Moreover, the number of plant food resources identified by *rbc*L (34 species) was higher than that by ITS2 (26 species) (S5 Table and S6 Table); however, ITS2 was used to supplement the plants that could not be identified by *rbcL* (Table 5). In this study, the number of identified plant food resources by using the combination of *rbcL* and ITS2 local databases (40 species) was higher than that obtained using the *rbcL* (34 species) or ITS2 (26 species) local database alone (Table 5).

**Table 7.** Comparison of the accuracy between this study and previous reports in estimating food resources using DNA metabarcoding.

| Scientific name | Analysis region | Presence or absence of local database | Species estimated accuracy (%) <sup>1)</sup> | References |
| --- | --- | --- | --- | --- |
| *Lagopus muta japonica* | *rbcL*+ITS2 | presence | 93.0 | This study |
| *Columba janthina nitens* | *trnL* intron (P6loop) | presence | 85.1 | [20] |
| *Anser fabalis* | *trnL* intron (P6loop) | presence | 75.0 | [26] |
| *Eonycteris spelaea* | *rbcL*+ITS2 | absence | 67.3 | [28] |
| *Anser albifrons* | *trnL* intron (P6loop) | presence | 66.7 | [26] |
| *Lemmus lemmus* | *trnL* intron (P6loop) | presence | 54.3 | [22] |
| *Microtus oeconomus* | *trnL* intron (P6loop) | presence | 52.9 | [21] |
| *Myodes rufocanus* | *trnL* intron (P6loop) | presence | 52.8 | [21] |
| *Apodemus speciosus* | *trnL* intron (P6loop) | absence | 43.3 | [29] |
| *Ctenomys minutus* | *trnL* intron (P6loop)+ITS1 | presence | 38.5 | [24] |
| *Apodemus argenteus* | *trnL* intron (P6loop) | absence | 35.5 | [29] |
| *Ctenomys flamarioni* | *trnL* intron (P6loop)+ITS1 | presence | 30.0 | [24] |
| *Pteropus hypomelanus* | *rbcL* | presence | 17.2 | [27] |

<sup>1)</sup> The species estimated accuracy was calculated by dividing the number of plant taxa of estimated to the species level by the number of total plant taxa.

Across all collection periods, the important plant foods for the Japanese rock ptarmigan in this study area were the family Ericaceae (98.1% of 105 fecal samples), especially *E. nigrum* var. *japonicum* (68.6%) as one of the dominant plant foods, which is consistent with the findings of a previous study conducted on Japanese rock ptarmigan elsewhere in Japan [13]. However, *V. ovalifolium* var. *ovalifolium* (69.5%) and *K. procumbens* (42.9%), which are frequently identified plant species of the Ericaceae family in this study, were not dominant species in the previous study [13]. These Ericaceae family plants were the dominant species in the community of alpine windswept dwarf shrubs of this study area. From May to June, Japanese rock ptarmigans forage mainly on alpine windswept dwarf shrubs, which have the earliest snowmelt in this study area. After July, the Japanese rock ptarmigans forage not only on alpine windswept dwarf shrubs but also on the community of snow-melted alpine leeward dwarf shrubs [1]. During our study, Japanese rock ptarmigans were found in both communities, but mainly foraged in alpine windswept dwarf shrubs. The dominant plant foods in the study area, such as *E. nigrum* var. *japonicum* (68.6%), and subdominant plants such as *P. aleutica* (13.3%), *V. vitis-idaea* (13.3%), and *B. ermanii* (11.4%) were mostly consistent with the finding of a previous study conducted on Japanese rock ptarmigan elsewhere in Japan [13], while other plant foods were endemic to this study area (Table 5). Notably, although *S. kurilensis* (14.3%), *N. crista-galli* subsp*. japonicum* (11.4%), and *P. juniperinum* (10.5%) are common species of other habitats elsewhere in Japan, there are no records of these species being utilized by Japanese rock ptarmigans in previous studies (to our knowledge) [13–15]. The reason for the absence of foraging records might be that the shoot height of these plants was lower than the Japanese rock ptarmigan’s eye level, so that the animal’s downward foraging behavior could not be adequately observed; however, the detailed reasons are unknown. The minor plant foods (less than 10% of 105 fecal samples) in this study area have not been reported previously [13–15]; plants with low populations within the study area (*e.g., Ixeridium dentatum* subsp. *kimuranum,* 7.6%; *Triantha japonica,* 5.7%), plants situated between masses of other plants (*e.g., Cornus canadensis,* 5.7%; *Gaultheria adenothrix,* 3.8%; *G. pyroloides,* 3.8%; *Coptis trifoliolata,* 1.9%), and plants that are difficult to identify from the external morphology (*e.g., Calamagrostis longiseta*, 6.7%; *Juniperus communis* var. *hondoensis*, 2.9%) can easily be overlooked in direct observation studies (Table 5). Moreover, hygrophytes (*e.g., N. crista-galli* subsp. *japonicum,* 11.4%; *J. filiformis,* 7.6%) growing in and around the small pools were identified for the first time in our study [13–15] (Table 5). This indicates that hygrophytes are also an important food resource for Japanese rock ptarmigans in the study area. In addition, *C. longiseta* (6.7%) in family Poaceae (Table 5), which is not a native plant of the alpine meadow zone and might have been imported by mountain climbers, was also an important plant food resource for Japanese rock ptarmigans in this study area.

On the other hand, *Arctous alpina* [14], *Arnica unalaschcensis* var. *tschonoskyi* [13], *Maianthemum dilatatum* [14], *Pedicularis chamissonis* var. *japonica* [13], and *Ranunculus acris* subsp. *nipponicus* [14] were also recorded as being foraged by other populations in previous studies [13–15] and grow in this study area; however, these plants were not detected in any of the fecal samples collected in this study (Table 5). The primer sets of *rbcL* and ITS2 used in this study are universal primers capable of amplifying *rbcL* and ITS2 in most plant species [33, 35]. In addition, both primer sets were able to amplify both rbcL and ITS of these plants (*A. alpina*, *P. chamissonis* var. *japonica*, *A. unalaschcensis* var. *tschonoskyi*, *M. dilatatum*, and *R. acris* subsp. *nipponicus*) in the construction of our local database, except for the ITS2 sequence of *M. dilatatum*.

Therefore, it is not expected that amplification of either *rbcL* or ITS2 will be affected by sequence variations in the regions corresponding to the primers in undetected plant species. Based on these considerations, it is reasonable to assume that these undetected plants species were not foraged by Japanese rock ptarmigans in this study area, but the reason for this is unknown. In addition, *Vaccinium shikokianum, V. smallii* var. *smallii*, and *V. uliginosum* var. *japonicum* were also recorded as forage plants of other populations in previous studies [13–15] and grow in this study area. We were unable to identify these three *Vaccinium* species at the species level in this study, as these three species have the same *rbcL* and ITS2 sequences. Therefore, it is necessary to select a region other than the *rbcL* and ITS2 regions for identification of such plant foods.

Of the subdominant plant foods (more than 10% to less than 30% of the 105 fecal samples), *S. commixta* (15.2%) and *S. kurilensis* (14.3%) were abundantly found in August (Fig 3 and Table 5). Since *S. commixta* is in full bloom and *S. kurilensis* is developing new leaves during this time, Japanese rock ptarmigans are thought to be selectively feeding on conspicuous flowers and soft new leaves. Since the DNA metabarcoding method cannot identify specific parts of the forage plants, direct observation during fecal sample collection can assist in clarifying the selectivity of foraging plant parts with phenological changes in alpine vegetation.

In general*, P. pumila* can be found in all Japanese rock ptarmigan habitats in Japan; however, individual habitats are characterized by their own unique alpine vegetation, *e.g, E. nigrum* var. *japonicum* is a dominant species in the study area of the Northern Japanese Alps, but rarely grows in the Japanese rock ptarmigan habitat of the Southern Japanese Alps [1]. The major plant food species for the Japanese rock ptarmigan may differ according to the mountain region [1]. Therefore, conservation of the Japanese rock ptarmigan requires us to clarify the relationship between the flora and plant food resources in different mountain systems, and to adapt vegetation conservation efforts to the flora of individual mountain regions.

## Conclusions

In this study area, the dominant taxa of the Ericaceae family in the alpine windswept dwarf shrub community were found to be the major plant foods for the Japanese rock ptarmigan from July to October, suggesting the need to manage this community to conserve their food resources. In addition, the local database constructed in this study can be used to survey other areas with similar flora. The Japanese rock ptarmigan does not fear people, allowing researchers to collect fecal samples efficiently and limiting the possibility of mistaking fecal samples for those of other birds. DNA barcoding using the local database is therefore considered to represent an important advance in our ability to elucidate the diet of the Japanese rock ptarmigan, and can also be used as a tool for conserving the alpine plant species that are their food sources.

## Acknowledgments

The authors thank the staff of the Taro-daira Koya and Ms. Chie Hashimoto, Ms. Haruka Kaga, Mr. Kouhei Tsuchimoto, Mr. Kouske Hujita, and Mr. Takeshi Shimamura of the College of Bioscience and Biotechnology at Chubu University for technical support.

## Supporting information

**S1 Table Results of homology search of each fecal sample using only the *rbcL* local database.** Bold type indicates an ASV of less than 98% homology.

**S2 Table Results of homology search of each fecal sample using only the ITS2 local database.** Bold type indicates an ASV of less than 98% homology.

**S3 Table Results of homology search using DDBJ of ASVs with less than 98% homology with the *rbcL* local.** Acceptances of more than 98% homology were adopted. Plant species that were unpublished and whose scientific names were not registered in BG Plants were excluded. In addition, we used data that were estimated to the species level.

**S4 Table Results of homology search using DDBJ of ASVs with less than 98% homology with the ITS2 local.** Acceptances of more than 98% homology were adopted. Plant species that were unpublished and whose scientific names were not registered in BG Plants were excluded. In addition, we used data that were estimated to the species level.

**S5 Table Results of food plant identification using *rbcL.*** 1) The plant species in parentheses are indicated by the upper taxonomic group because they had the same DNA sequence.

**S6 Table Results of food plant identification using ITS2.** 1) The plant species in parentheses are indicated by the upper taxonomic group because they had the same DNA sequence.

